# Neuroprotective Effects of *Nigella sativa* Extract in a Rat model of Propionic Acid-Induced Autism Spectrum Disorder

**DOI:** 10.1101/2025.08.26.672514

**Authors:** Sultan M. Alshahrani, Sultan F. Kadasah, Humara Jan, Farid Menaa

## Abstract

**Background:** Autism Spectrum Disorder (ASD) is neurodevelopmental disorder with characteristics of impairments in social interaction, communication, and repetitive behaviors. Current pharmacological interventions often focus on alleviating symptoms, but they rarely address the underlying neurobiological disruptions or have sustained therapeutic effects.

**Aims:** The present study explores the therapeutic effect of Nigella sativa (black cumin) extract in a propionic acid (PPA)-induced wistar rat model of ASD.

**Methods:** Forty male Wistar rats (n = 10 per group) were randomly assigned to four experimental groups, including control, PPA, N. sativa Low-Dose (NSL), and N. sativa High-Dose (NSH). The control group received daily intravenous (i.v.) injections (1 mL/kg) of saline (0.9% NaCl) over five days. The PPA group received daily intraperitoneal (i.p.) injections of 250 mg/Kg PPA for five days to induce ASD-like behaviors. NSL and NSH groups received N. sativa extract (10 mg/Kg and 50 mg/Kg, respectively) orally for 28 days, starting 7 days before PPA administration. Behavioral assessments, including social interaction, stereotypic behaviors, and anxiety, were conducted on day 28. Then, biochemical analyses (i.e., oxidative stress, inflammatory markers, neurotransmitter levels, caspase-3 expression, and histopathology) were performed.

**Results:** The PPA group showed significantly (p < 0.05) reduced social interaction and increased stereotypic behaviors and heightened anxiety-like responses, indicative of ASD-like symptoms. Treatment with *N. sativa* resulted in a significant (p < 0.05) improvement in social behaviors, a reduction in stereotypic behaviors, and a decrease in anxiety-like responses. Biochemical analysis revealed that N. sativa treatment significantly (p < 0.05) reduced oxidative stress, evidenced by lower MDA levels, higher GSH, and restored SOD activity. In both NSL and NSH groups, it was observed that (i) inflammatory cytokines (i.e., TNF-α and IL-1β) were significantly (p < 0.05), (ii) neurotransmitter levels (i.e., dopamine and serotonin) were normalized, (iii) caspase-3 expression was significantly reduced, leading to a reduction in neuronal apoptosis-induced cell death, and (iv) histopathological analysis revealed reduced neuronal damages and a glial activation.

**Conclusions:** *N. sativa* extract appears highly effective in ameliorating ASD behavioral and biochemical pathology abnormalities associated with ASD in a PPA-rat model. These results underscore the role of *N. sativa for* the treatment of ASD, including as possible adjunctive therapeutic option for this challenging neurodevelopmental disorder, though further research studies are necessary, including to determine the key phytochemical(s) involved in such beneficial effects.

**Highlights:**

- High-dose *Nigella sativa* (NSH) acted as a potent brain antioxidant and anti-inflammatory agent;
- NSH restored neurotransmitter balance by normalizing dopamine and serotonin levels;
- NSH significantly reduced propionic acid (PPA)-induced neuronal apoptosis and caspase-3 expression;
- NSH minimized neuronal damages and preserved brain tissue structure;
- NSH improved core ASD-like behaviors, including social deficits and stereotypy.

## I. Introduction

Autism Spectrum Disorder (ASD) is a multifaceted neurodevelopmental disorder which affects about one in 54 children worldwide and shows higher prevalence in men [1]. It is defined by impairments in social communication and interaction, restricted and repetitive behaviors, as well as sensory sensitivities [2]. The presenting features of ASD are highly heterogeneous, encompassing individuals with mild ASD symptoms to those with markedly impaired features that profoundly affect functional living [3]. Although ASD is very common, the precise causes are still unknown, but the underlying aetiology seems to involve a mixed genetic, environmental and neurobiological background [4].

Current treatment strategies primarily focus on behavioural therapies, speech and language interventions, and pharmacological approaches aimed at managing specific symptoms such as anxiety, irritability, or hyperactivity. Nevertheless, there is no single treatment that has been universally successful and existing pharmacological treatments are often accompanied by adverse effects and/or lack of ability to target underlying neurobiological reasons for the disorder [5]. This has sparked an increasing interest in the research of alternative therapeutic approaches, especially those which are safe, affordable and can normalize neurological and behavioural symptoms of ASD [6].

One promising area of research lies in the use of traditional herbal medicines, which has gained cumulative popularity in today’s medical practice. Indeed, herbal medicines are used to manage various ailments, and some of them (like *N. sativa*) exert potential neuroprotective and therapeutic effects in neurodevelopmental disorders like ASD [7, 8].

Herbal medicines are considered as promising adjuvants for ASD, most likely because of their natural properties, mild side-effect profiles, and availability [9]. Many plants exert antioxidant, anti-inflammatory, and neuroprotective activities that are relevant for the treatment of ASD symptoms [10].

*N. sativa* (Kalaunji, also known as black cumin or black cumin), a versatile spice often used in traditional Indianm Middle Eastern and African dishes, has emerged as a highly promising agent for neurological protection and therapeutic uses [11]. Its seeds, which contain a variety of bioactive compounds such as thymoquinone, nigellidine, and other alkaloids, have been studied for their anti-inflammatory, antioxidant, and immune-modulatory effects [12, 13]. Recent works have indicated Nigella sativa as a potential neuroprotective agent that could decrease neuroinflammation, improve neuronal function, and prevent oxidative stress which are thought to be of important contributory roles in ASD formation and progression [14].

Oxidative stress and neuroinflammation have consistently been reported in the brain of ASD. Higher concentrations of reactive oxygen species (ROS), pro-inflammatory cytokines and imbalance between antioxidant defence and oxidative stress are frequently found in ASD patients [15]. These alterations are presumed to play a role in neurodevelopmental delays and cognitive deficits [15]. Animal models of ASD are essential for understanding its complex neurobiology and testing potential therapies. Rodent models are widely used as they can mimic core ASD-like features such as impaired social interaction, increased anxiety, and stereotypic behaviors [16]. These models are typically generated through genetic manipulation or environmental exposures, including prenatal infections, valproic acid (VPA), or propionic acid (PPA) administration [17]. Among these, chemically induced models like PPA are particularly valuable as they allow for the induction of ASD-like symptoms postnatally and reflect gut-brain axis involvement, a key emerging mechanism in ASD [18]. Such models provide reproducible behavioral and biochemical alterations and are widely used to evaluate natural or pharmacological interventions for ASD.

Considering the potent antioxidant and anti-inflammatory activities of *N. sativa* [12, 13], its possible ability to reduce the above pathological features gives N. sativa the potential ability to be explored as a treatment for ASD. To the best of our knowledgem and given the growing evidence supporting the therapeutic potential of *N. sativa* in neurological conditions, this study aims, for the first time, to evaluate its protective effects in a rat model of ASD induced by propionic acid (PPA). Thereby, behavioral, biochemical and histological investigated have been carefully conducted to determine whether *N. sativa* is able to ameliorate ASD-like symptoms induced by PPA adminitration.

## II. MATERIALS and Methods

### 2.1. Preparation of *Nigella sativa* extract

*N. sativa* seeds were purchased locally and their identity was verified at our department by an experimented herborist. The seeds were washed, dried, and then crushed into a fine powder. A total of 20 g of the powdered seeds was added to 400 mL of distilled water, and extraction was carried out using steam distillation. The yield of the extract was 0.5%, determined as a percentage of dry weight of the seeds. The extract was stored in screw-capped tubes, protected from light, and kept at 20°C until use.

### 2.2. Animals and ethical consideration

Forty male Wistar rats, aged 8 to 10 weeks and weighing between 200 and 250 g, were purchased from a certified animal supplier. The animals were housed under standard laboratory conditions (i.e., 12-hour light/dark cycle, temperature of 22 ± 2°C, humidity of 50–60%), and to minimize the stress, the rats were allowed to acclimatize to their environment for 7 days before the start of the experiment [19]. During the acclimatization period, the rats had *ad libitum* access to standard rodent chow and filtered water. Rats were constantly monitored for distress or disease.

Ethical approval for the study was obtained from the Institutional Animal Care and Use Committee (IACUC), and all procedures adhered to national and institutional animal welfare guidelines.

### 2.3. Experimental design

Rats (n=40) were assigned at random to four experimental groups of 10 animals each. Group I (control) received daily intravenous (i.v.) injections (1 mL/kg) of saline (0.9% NaCl) over five days, to obtain baseline values for normal behavior and physiology; Group II (PPA) received propionic acid *via* intraperitoneal (i.p.) route at a dose of 250 mg/kg/day for five days [20], to produce ASD-like behavioural changes; Group III (NSL) received low-dose of N. sativa extract in *per os* (p.o.; orally) at a dose of 10 mg/kg/day for 28 days, starting 7 days prior to the PPA injections; Group IV (NSH) received low-dose of *N. sativa* extract in *per os* (p.o.; orally) at a dose of 50 mg/kg/day for 28 days, starting 7 days prior to the PPA injections; this higher dose aimed to determine whether it would have a more significant effect in modulating PPA-induced ASD-like symptoms and improving neurodevelopmental outcomes. The dosage was selected based on previous studies [21].

After 28 days of treatment, the animals underwent behavioral testing to assess social interactions, anxiety, stereotypic behaviors, and motor activity, which are typically associated with ASD. After the behavioral tests, the rats were humanely euthanized using an overdose of sodium pentobarbital (≥ 150 mg/kg, intraperitoneally (i.p.)), a method recommended to minimize pain and distress, in accordance with the guidelines of the Institutional Animal Care and Use Committee (IACUC). Subsequently, their brain and peripheral tissues were extracted for histological, immunohistochemical and biochemical assays.

#### 2.4.1. Behavioral testing

Behavioral tests were conducted at day 28 in order to assess social behavior, anxiety, and stereotypic behaviors that are characteristic of ASD. All tests were performed during the light cycle in a controlled environment, and the order of testing was randomized [22].

The *social behavior* was measured through the reciprocal social interaction test [22], particularly the ability to interact with an unfamiliar conspecific. Each experimental rat was placed in a neutral open-field arena (60 cm × 60 cm) along with an age- and sex-matched conspecific that was not part of the treatment groups and had no prior exposure to behavioral testing (n = 40). Rats were introduced into a neutral arena (60 cm x 60 cm) with an age- and sex-matched rat. The test rat was given free access to the apparatus for 10 min. Interactions including sniffing, following, and grooming were were observed and recorded manually.. Physical contact time and total number of interactions were compared.

The *anxiety* disorder was determined through the open field test (OFT) and elevated plus maze test (EPMT). For the OFT, box arena (50 cm x 50 cm x 30 cm) with completely black sides and floor composed of 25 equal tiles was employed to evaluate overall locomotor activity and anxiety-like/avoidant behaviors. Rats were confined to the centre of the arena and free to roam for 10 min. Total travelled distance as well as the time spent in the centre of the arena were quantified. Decreased activity and fewer centre minutes were interpreted as being correlates for a higher level of anxiety. For the EPMT, the maze consisted of two open arms (50 cm × 10 cm) and two enclosed arms (50 cm × 10 cm), elevated 50 cm above the ground. Every rat was placed in the middle and free to explore for 5 min. Time spent in the open *versus* enclosed arms were recorded. The greater was the time spent in the confined arms, the greater anxiety was [23].

The *stereotypic behavior* tests (e.g., like repetitive grooming, pacing, and head bobbing) indicated ASD-related traits, and consisted in observing the rats for 10 minutes in their clean home cages. The frequency, duration, and intensity of the behaviors were recorded after a 10 min acclimation period [24].

#### 2.4.2. Biochemical assays

Following the behavior tests, *ex-vivo* biochemical assays were conducted in the brain of euthanized rats to assess the concentration levels of oxidative stress and pro-inflammation markers, as well as of key neurotransmitters.

Brain-specific oxidative stress in the brain was evaluated by thiobarbituric acid-reactive substances (TBARS) as the indicator of lipid peroxidation based on MDA levels determined following the method of Placer et al. [25]. Additionally, glutathione (GSH) levels were measured using Ellman’s method [26], and superoxide dismutase (SOD) activity was assessed according to the method of Marklund et al. [27].

The levels of pro-inflammatory cytokines in brain tissue, including tumor necrosis factor-alpha (TNF-α) and interleukin-1 beta (IL-1β), were measured using ELISA kits from RayBiotech (Peachtree Corners, GA, USA), while the levels of neurotransmitters such as dopamine and serotonin were assessed using ELISA kits from Invitrogen (Thermo Fisher Scientific, Waltham, MA, USA). All assays were performed according to the respective manufacturers’ protocols, and absorbance was recorded at 450 nm using a microplate reader [28].

#### 2.4.3. Immunochemistry

To assess apoptosis in the brain, immunohistochemical staining for caspase-3 was performed. After euthanizing the animals, brain sections were obtained and fixed in 10% formalin. After paraffin embedding, 5 µm-thick sections were prepared [19] and underwent antigen retrieval using citrate buffer. Endogenous peroxidase activity was blocked with 3% hydrogen peroxide (H_2_O_2_) for 30 minutes. Sections were incubated overnight at 4°C with a primary antibody against caspase-3 (1: 200 dilution; Cell Signaling Technology, Danvers, Massachusetts, USA). A biotinylated secondary antibody was applied and subsequently visualized by diaminobenzidine (DAB). Sections were counterstained with haematoxylin, and the number of caspase-3 positive cells were counted manually in five randomly selected high-power fields (400×) under a light microscope.

#### 2.4.4. Histopathology

Brain tissues were stained with haematoxylin and eosin (H&E) to determine general morphology and tissue architecture [19]. After fixation in formalin and paraffin embedding, 5 µm-thick sections were cut and stained with H&E. The stained slices were observed by using a light microscope (Olympus BX53, Olympus Corporation, Tokyo, Japan). to evaluate the structural abnormalities such as neuronal degeneration, glial cell activation, and brain structural changes suggestive of neurodevelopmental changes in response to the PPA-induced ASD. Histopathological changes were scored based on standardized criteria.

### 2.5. Statistical analysis

Data are shown as mean ± standard error (SE). Statistical analyses were performed using SPSS version 11 (SPSS Inc., Chicago, IL, USA). A one-way analysis of variance (ANOVA) was used to compare between-group differences, followed by Duncan’s post hoc tests to identify specific group differences. Statistical significance was set at p < 0.05.

## III. Results

### 3.1. Effect of *N. sativa* extract on social interaction and stereotypic behaviors of ASD-like rats

In the reciprocal social interaction **(Fig. 1A)** and stereotypic behavior **(Fig. 1B)** tests, significant (p < 0.05) differences were observed between the experimental groups of ASD-like Wistar rats.

**Figure 1.**
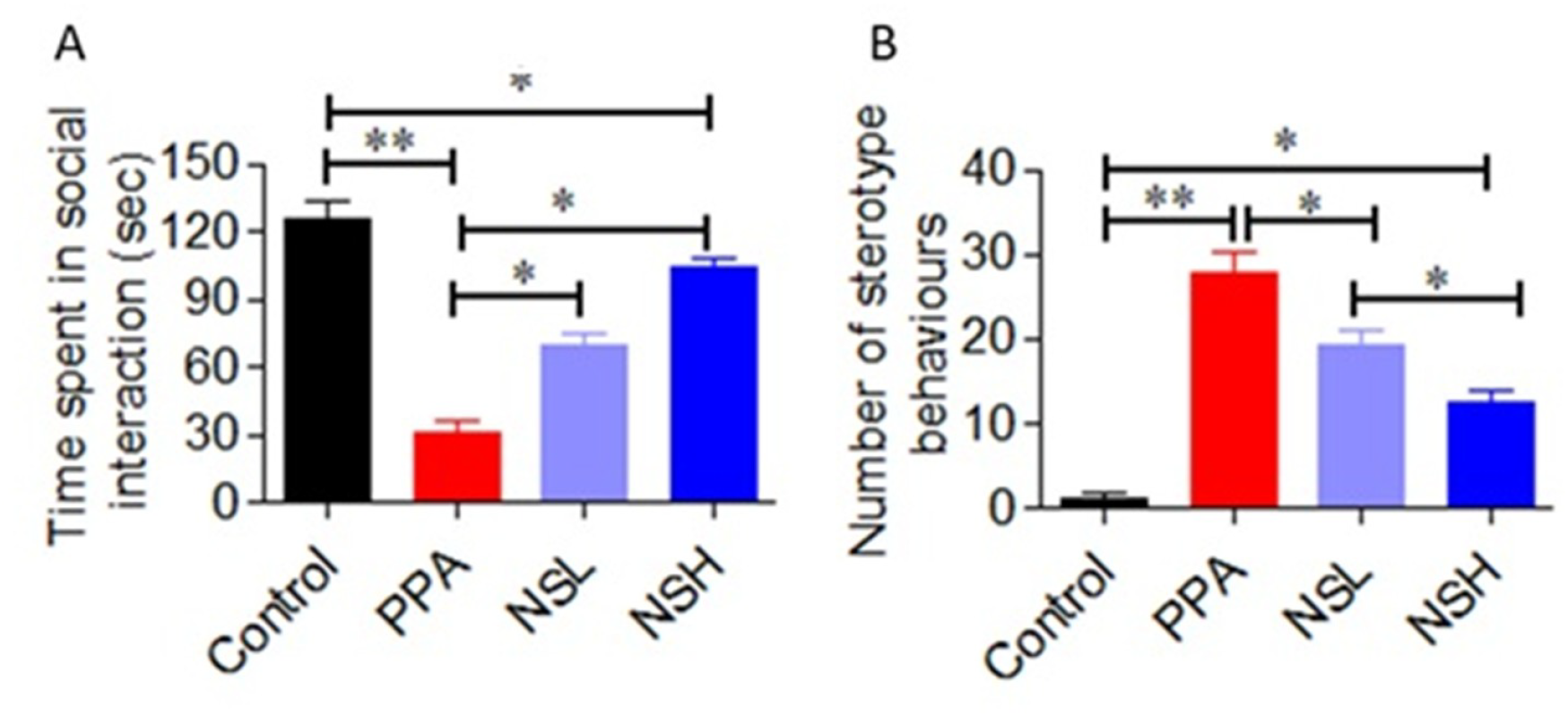
Effect of *Nigella sativa* extract on social interaction and stereotypic behaviors in autism-spectrum disorder (ASD)-like adult Wistar rats (N=40, n=10/group). **(A)** Time spent in (reciprocal) social interaction; **(B)** Number of stereotype behaviors (i.e., repetitive grooming, pacing, and head bobbing). Control (Group I, NaCl 0.9%); PPA (Group II, Propionic acid); NSL (Group III, *N. sativa* Low-dose); NSH (Group IV, *N. sativa* High-dose). Data are presented as mean ± SD; *p < 0.05, **p < 0.005.

The Group I (control, NaCl) exhibited high levels of social interaction (125.5 ± 23.6 seconds) and negligible stereotypic behaviors (1.30 ± 0.34). In comparison, the Group II (PPA-induced ‘ASD’, abreviated as PPA) demonstrated a marked reduction in social interactions (31.5 ± 15.13 seconds) and a significant increase in stereotypic behaviors (27.9 ± 8.03 seconds), suggestive of ASD-like traits. Both *N*.*sativa* treatment groups III (NSL) & IV (NSH) showed significant improvements in social behavior compared to Group II. Also, the NSH Group demonstrated the most substantial recovery (104.5 ± 12.4 seconds), compared to the NSL (69.9 ± 17.8 seconds, with social interaction frequency and time spent with the unfamiliar rat approaching control levels. In the stereotypic behavior test, the NSH Group showed the most pronounced reduction (p < 0.05) in repetitive behaviors (12.4 ± 4.7 seconds), compared to the NSL Group (19.4 ± 5.3 seconds), strongly suggesting a dose-dependent response.

### 3.2. Effect of *N. sativa* extract on anxiety-like behaviors of ASD-like rats

In the Elevated Plus Maze Test (EPMT), the time spent in the closed arms **(Fig. 2A)** was significantly altered across groups. The Control Group exhibited a mean closed arm time of 145.9 ± 17.4 sec, reflecting balanced exploratory behaviour. In contrast, the PPA Group showed a marked increase in closed arm time (243.1 ± 25.8 sec, *p* < 0.05 vs. Control), indicating heightened anxiety-like behaviour. Treatment with *Nigella sativa* led to a dose-dependent improvement. The NSL Group recorded a moderate reduction in closed arm time (189.7 ± 24.1 sec, *p* < 0.05 vs. PPA), while the NSH Group demonstrated the most pronounced anxiolytic effect with a significantly reduced closed arm time (154.5 ± 18.6 sec, *p* < 0.05 vs. PPA), approaching values observed in the Control Group.

**Figure 2.**
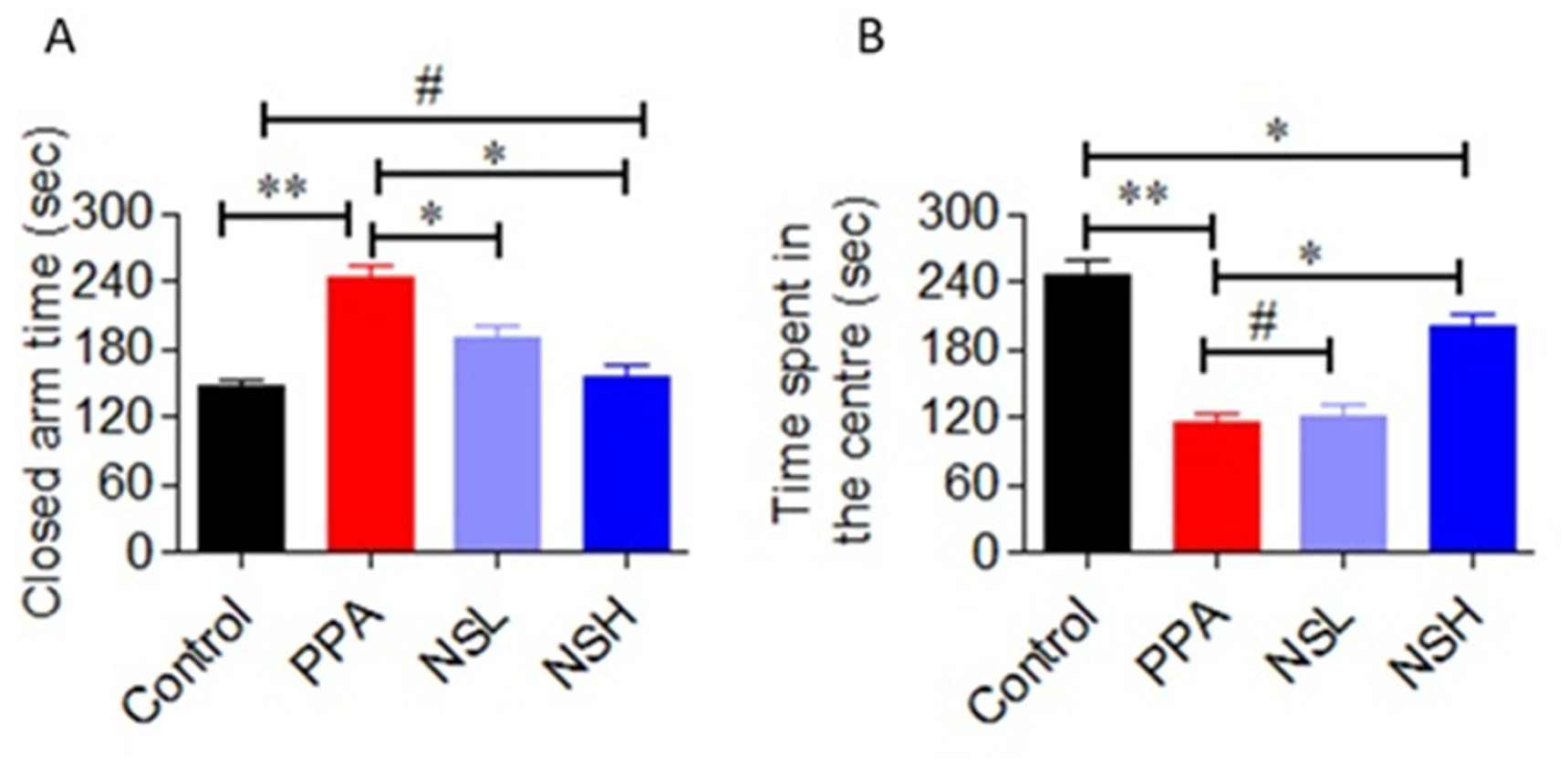
Effect of *Nigella sativa* extract on anxiety-like behaviors in autism-spectrum disorder (ASD)-like adult Wistar rats (N=40, n=10/group). **(A)** Closed arm time; **(B)** Time spent in the centre in control. Control (Group I, NaCl 0.9%); PPA (Group II, Propionic acid); NSL (Group III, *N. sativa* Low-dose); NSH (Group IV, *N. sativa* High-dose). Data are presented as mean ± SD; *p < 0.05, **p < 0.005, ^#^p > 0.05 (ns).

In the OFT, the time spent in the centre of the arena **(Fig. 2B)** was significantly reduced in the PPA Group (114.5 ± 28.9 sec) compared to the Control Group (244.6 ± 31.8 sec). Both treatment groups demonstrated a significant improvement: NSL rats spent 120.5 ± 10.7 sec in the centre, while NSH rats reached 201.6 ± 22.1 sec, significantly higher than PPA (p < 0.05). In parallel, the total distance travelled in the OFT **(Suppl. File, Fig. S1A)** was markedly lower in the PPA Group (3470 ± 415 cm) than in the Control Group (5249 ± 447 cm). The NSL Group travelled 4939 ± 530 cm, while the NSH Group showed a significant increase (5707 ± 477 cm, p < 0.05 vs. PPA), reflecting restoration of normal locomotor activity.

Furthermore, the time spent in the open arms of the EPM **(Suppl. File, Fig. S1B)** was drastically reduced in the PPA Group (45.6 ± 11.5 sec) relative to Controls (126.7 ± 20.8 sec). The NSL and NSH Groups exhibited a progressive recovery, spending 77.9 ± 16.3 sec and 103.3 ± 22.1 sec, respectively, in the open arms (p < 0.05 vs. PPA), indicating decreased anxiety-like behaviour. These findings indicate that *N. sativa* extract effectively mitigated anxiety-like behaviour in ASD-like rats, particularly at higher doses, as evidenced by the reduced preference for enclosed arms in the EPMT.

### 3.3. Effect of *N. sativa* extract on oxidative stress and inflammatory markers in ASD-like rats

The expression levels of oxidative stress have been evaluated using ELISA method. In case of oxidative stress, malondialdehyde (MDA) **(Fig. 3A)**, superoxide dismutase (SOD) **(Fig. 3B)**, and glutathione (GSH) **(Fig. 3C)** activities were quantified from the brain of ASD-like rats, using routinely lab-used methods.

**Figure 3.**
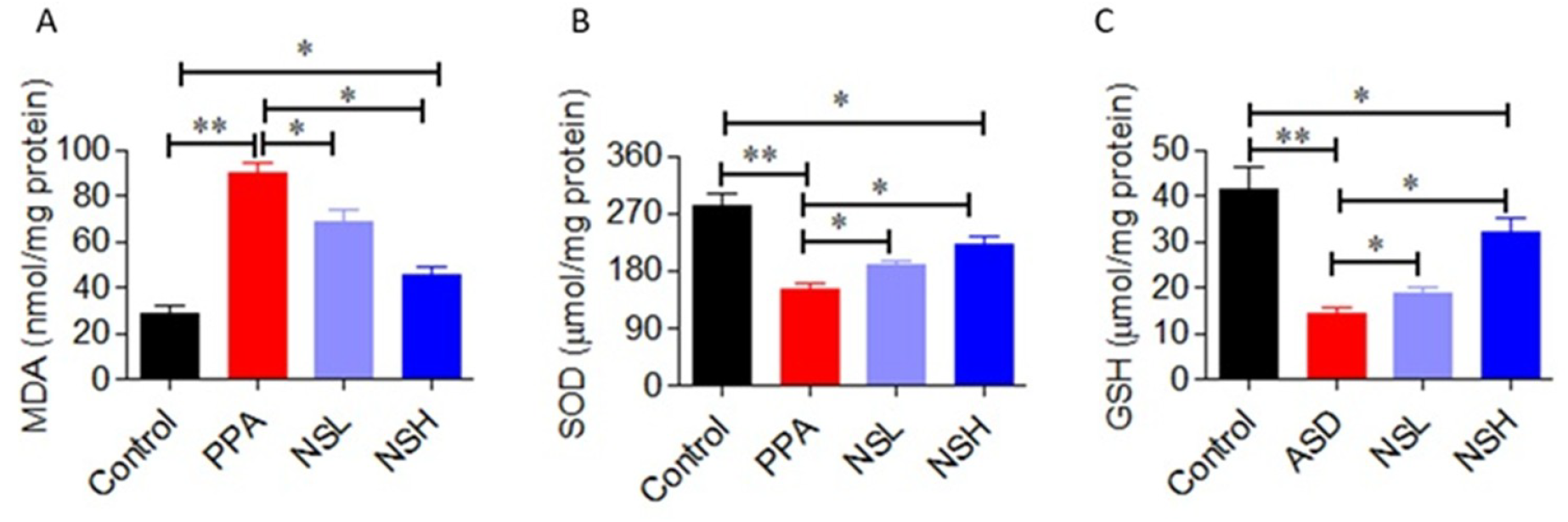
Effect of *Nigella sativa* extract on oxidative stress markers in autism-spectrum disorder (ASD)-like adult Wistar rats (N=40, n=10/group). **(A)** malondialdehyde (MDA); **(B)** superoxide dismutase (SOD); **(C)** glutathione (GSH). Control (Group I, NaCl 0.9%); PPA (Group II, Propionic acid); NSL (Group III, *N. sativa* Low-dose); NSH (Group IV, *N. sativa* High-dose). Data are presented as mean ± SD; *p < 0.05, **p < 0.005.

Comparatively to Control Group in which lowest levels of MDA (28.5 ±2.2 nmol/mg protein) as well as highest levels of GSH (41.2 ±3.4 µmol/mg protein) and SOD (283.1±14.5 µmol/mg protein) activities were noticed, the PPA Group showed significant (p < 0.05) oxidative stress, as evidenced by elevated levels of MDA (89.8 ± 15.2 nmol/mg protein) as well as reduced levels of SOD (151.5 ± 30.3 µmol/mg protein) and GSH (14.4 ± 3.2 µmol/mg protein), indicating increased lipid peroxidation and diminished antioxidant defence.

Comparatively to the PPA Group, the treatment with *N. sativa* resulted in a dose-dependent improvement of these oxidative stress markers; these beneficial effects were partial but significant (p < 0.05). Indeed, the NSL Group displayed better antioxidative stress than the PPA Group (MDA: 68.6 ± 9.2 nmol/mg protein; SOD: 188.2 ± 8.3 µmol/mg protein; GSH: 18.9 ± 4.2 µmol/mg protein); interestingly, the NSH Group exhibited the most significant improvements, with drastic reduction of MDA levels (45.6 ± 10.1 nmol/mg protein) and augmentation of SOD (222.7 ± 36.1 µmol/mg protein) and GSH (32.2 ± 9.08 µmol/mg protein) levels, indicating a capacity of such a concentrayion of *N. sativa* extract (50 mg/kg/day) in reverting substantially (almost to control levels) the PPA-induced oxidative stress in ASD-like rats.

### 3.4. Effect of *N. sativa* extract on neuroinflammatory markers in ASD-like rats

In case of pro-inflammatory markers, the expression levels of tumor necrosis factor-alpha (TNF-α) **(Fig. 4A)** and interleukin-1 beta (IL-1β) **(Fig. 4B)** were assessed by ELISA method from the brain of ASD-like rats.

**Figure 4.**
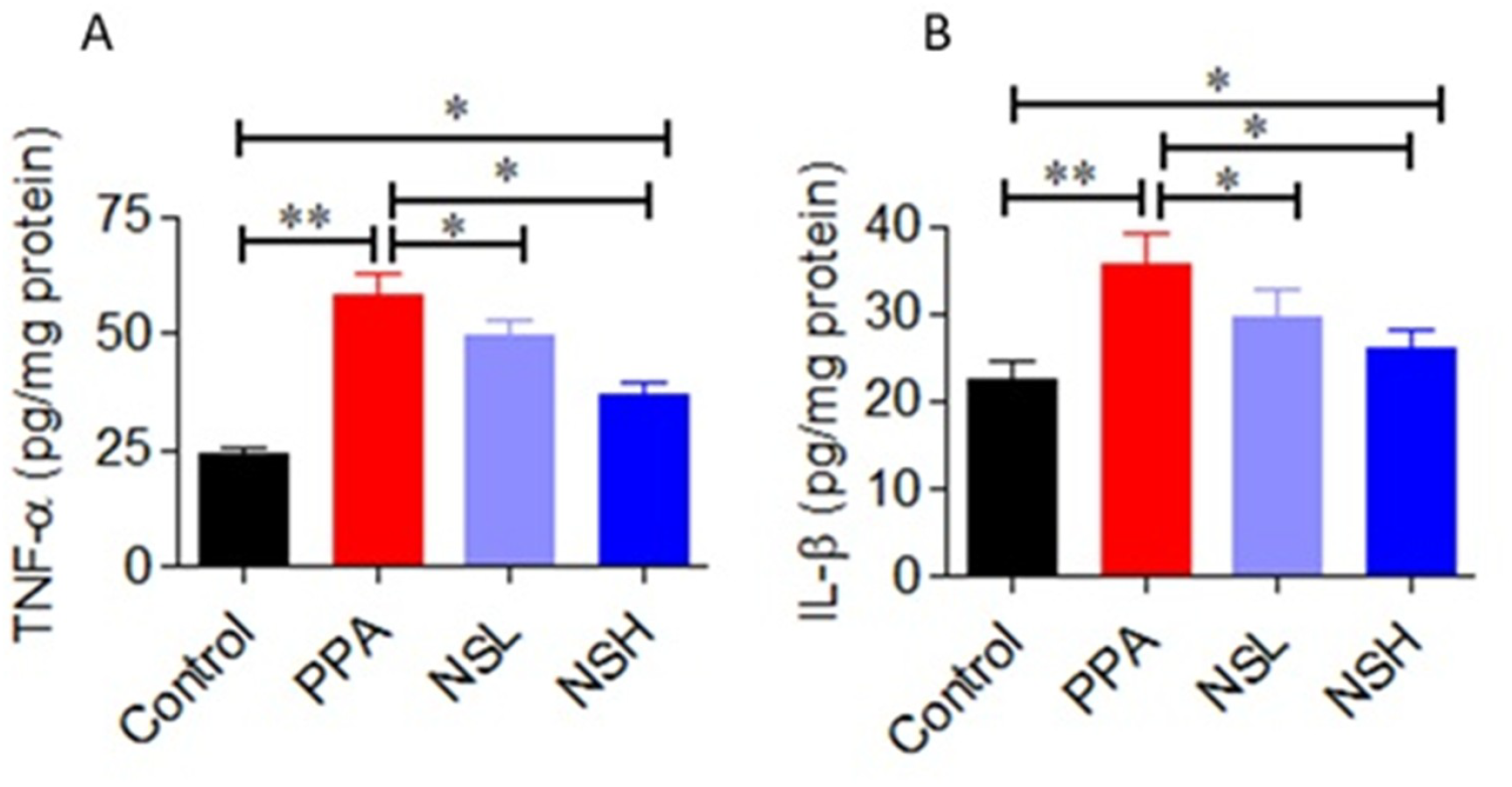
Effect of *Nigella sativa* extract on pro-inflammatory cytokines in autism-spectrum disorder (ASD)-like adult Wistar rats (N=40, n=10/group). **(A)** tumor necrosis factor-alpha (TNF-α); **(B)** interleukin-1 beta (IL-1β). Control (Group I, NaCl 0.9%); PPA (Group II, Propionic acid); NSL (Group III, *N. sativa* Low-dose); NSH (Group IV, *N. sativa* High-dose). Data are presented as mean ± SD; *p < 0.05, **p < 0.005.

The Control Group shows TNF-α and IL-1β levels of 24.0 ± 2.2 pg/mg protein and 22.5 ± 4.8 pg/mg protein, respectively. Comparatively to this Group, the PPA Group significantly (p < 0.05) elevated the levels of pro-inflammatory cytokines TNF-α (57.8 ± 13.02 pg/mg protein) and IL-1β (35.5 ± 12.1 pg/mg protein).

Both *N. sativa*-treated groups (NSL and NSH) demonstrated significant (p < 0.05) reductions of these pro-inflammatory markers, with the NSL exhibiting (TNF-α: 49.4 ± 3.2 pg/mg protein; IL-1β: 29.5 ± 4.1 pg/mg protein), and the NSH Group exhibiting the greatest anti-inflammatory effects (TNF-α: 37.1 ± 7.8 pg/mg protein; IL-1β: 26.1 ± 6.08 pg/mg protein), strongly suggesting a dose-dependent response of the phytoextract in ASD-like rats.

### 3.5. Effect of *N. sativa* extract on neurotransmitters in ASD-like rats

The expression levels of the neurotransmitters dopamine **(Fig. 5A)** and serotonin (IL-1β) **(Fig. 5B)** were assessed by ELISA method from the brain of ASD-like rats.

**Figure 5.**
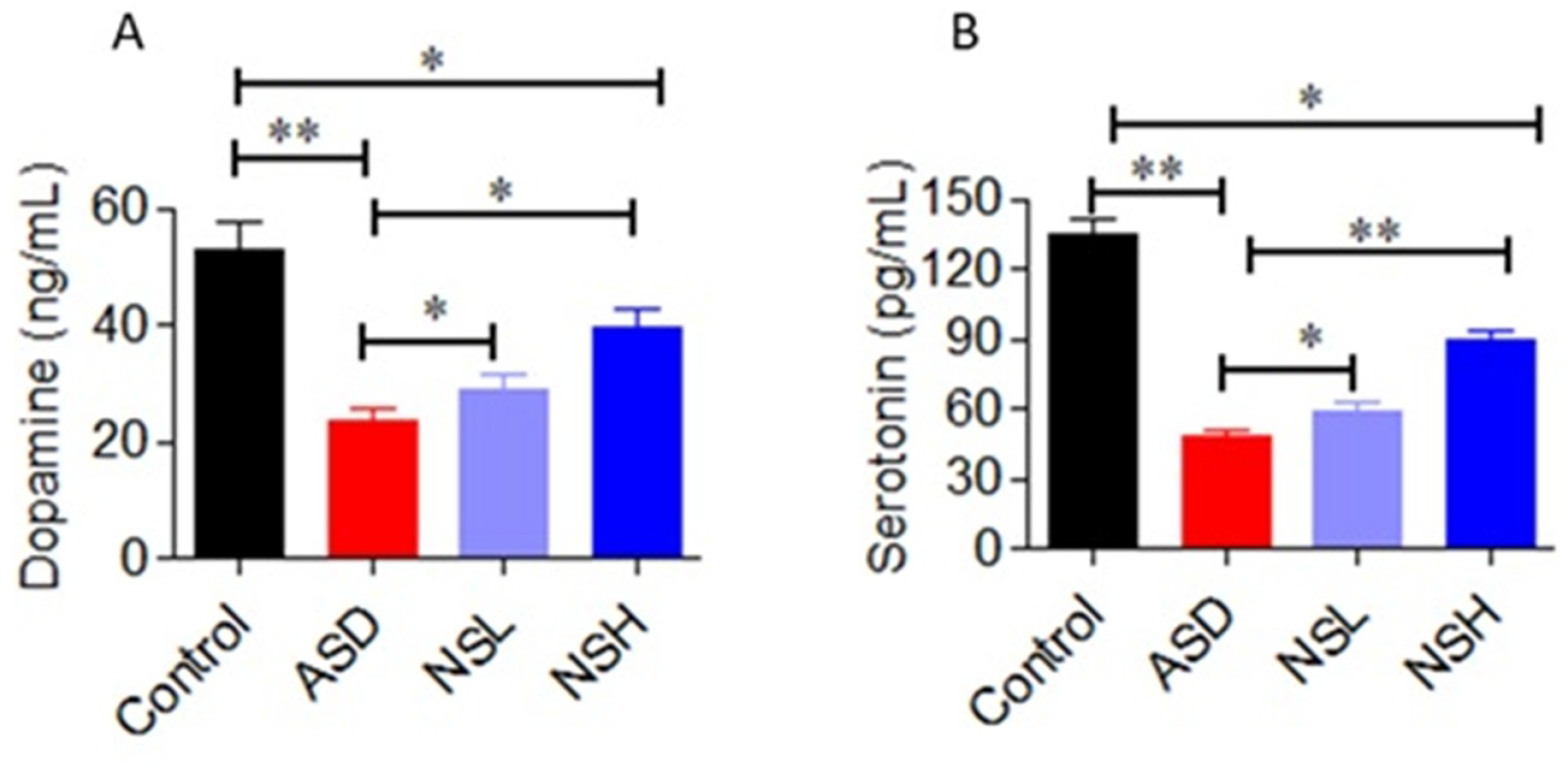
Effect of *Nigella sativa* extract on neurotransmitters in autism-spectrum disorder (ASD)-like adult Wistar rats (N=40, n=10/group). **(A)** dopamine; **(B)** serotonine. Control (Group I, NaCl 0.9%); PPA (Group II, Propionic acid); NSL (Group III, *N. sativa* Low-dose); NSH (Group IV, *N. sativa* High-dose). Data are presented as mean ± SD; *p < 0.05, **p < 0.005.

Comparatively to the Control Group (dopamine: 52.7 ng/mL; serotonin: 134.9 pg/mL), the PPA Group exhibited significant dysregulation in neurotransmitter levels, with decreased concentrations of dopamine (23.8 ± 6.19 ng/mL) and serotonin (48.1 ± 9.9 pg/mL).

Both *N. sativa*-treated groups (NSL and NSH) showed dose-dependent restoration of neurotransmitter balance. Indeed, NSL Group showed improvements in both neurtransmitters expression levels (dopamine: 29.1 ± 1.8 ng/mL; serotonin: 58.3 ± 4.5 pg/mL), with the NSH Group exhibiting the most significant (p < 0.05) improvement (dopamine: 39.7 ± 10.3 ng/mL; derotonin: 89.1 ± 14.07 pg/mL).

### 3.6. Effect of *N. sativa* extract on neuroapoptosis in ASD-like rats

The neuronal cell apoptosis was assessed in triplicate by immunochemistry of caspase-3 from the brain of euthanized ASD-like rats **(Fig. 6)**. The apoptotic cells turned brown after staining and were counted in at least three independent fields and slides.

**Figure 6.**
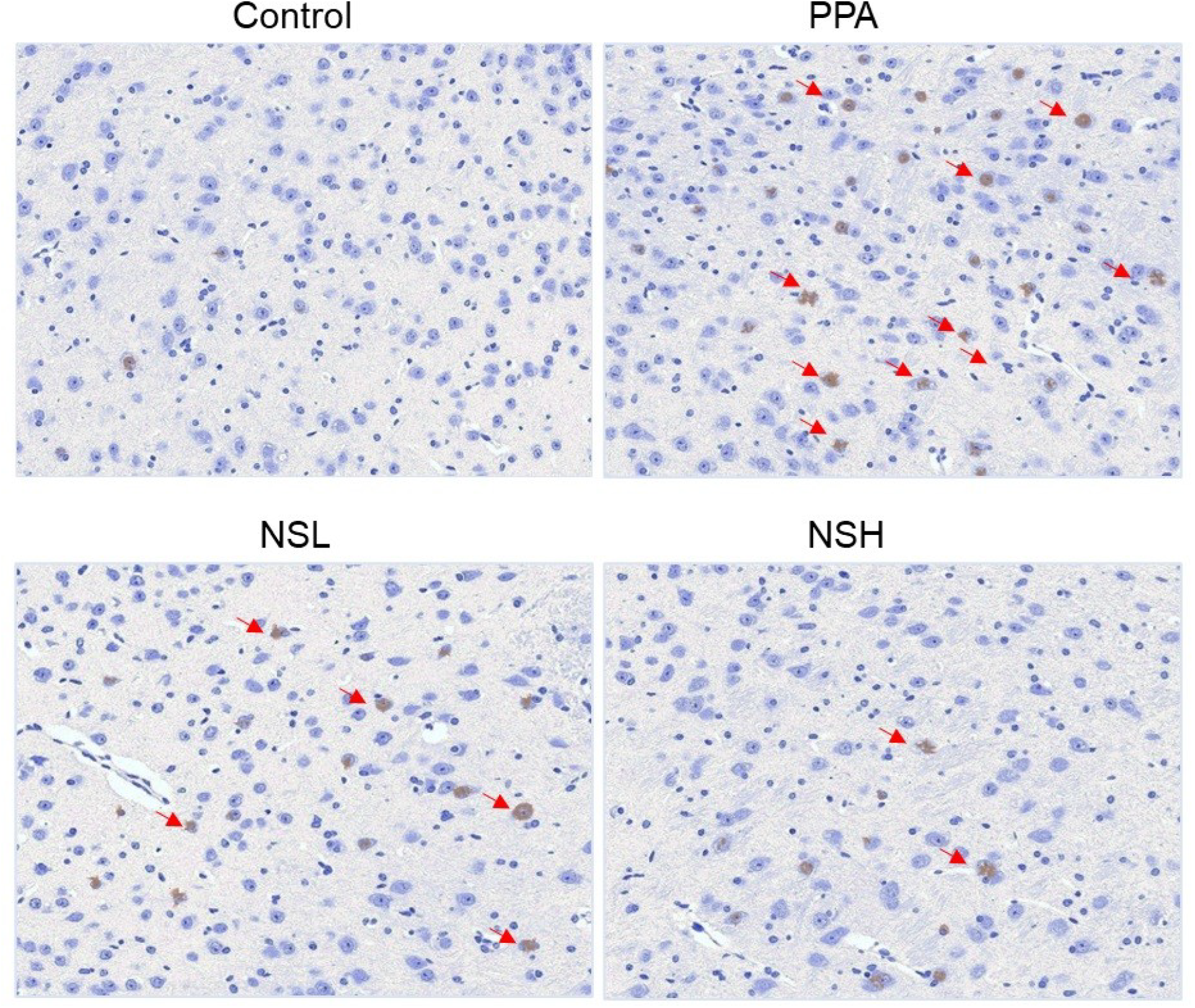
Effect of *Nigella sativa* extract on caspase-3 expression in autism-spectrum disorder (ASD)-like adult Wistar rats (N=40, n=10/group). Control (Group I, NaCl 0.9%); PPA (Group II, Propionic acid); NSL (Group III, *N. sativa* Low-dose); NSH (Group IV, *N. sativa* High-dose). Red arrows depict caspase-3 expression.

Comparatively to the Control Group (2 ± 1 apoptotic/brown-stained cells), the brain tissue section of PPA-treated rats showed significantly higher (p < 0.05) caspase-3 expression in cells (45 ± 5 brown-stained cells), indicating increased neuronal cell death by apoptosis.

In both *N. sativa*-treated groups (NSL and NSH), micrographs depicted significant reductions (p < 0.05 compared to PPA Group) in caspase-3 expression levels (about 3 folds), with a number of apoptotic neuronal cells in the NSL Group of 15 ± 3, and with the NSH Group exhibiting the most pronounced anti-apoptotic effects with 9 ± 2 brown-stained cells.

### 3.7. Histopathological effect of *N. sativa* extract on brain tissue of ASD-like rats

Histopathological examination of brain tissue of ASD-like rats **(Fig. 7)** was performed in triplicate from three independent fields and slides. The brain tissue architecture of rats used as Control was ‘normal’ and did reveal a negligeable number of alterations. However, in the PPA group, a high number of neuronal damages (11 ± 4) can be noticed; these changes are consistent with neurodevelopmental alterations.

**Figure 7.**
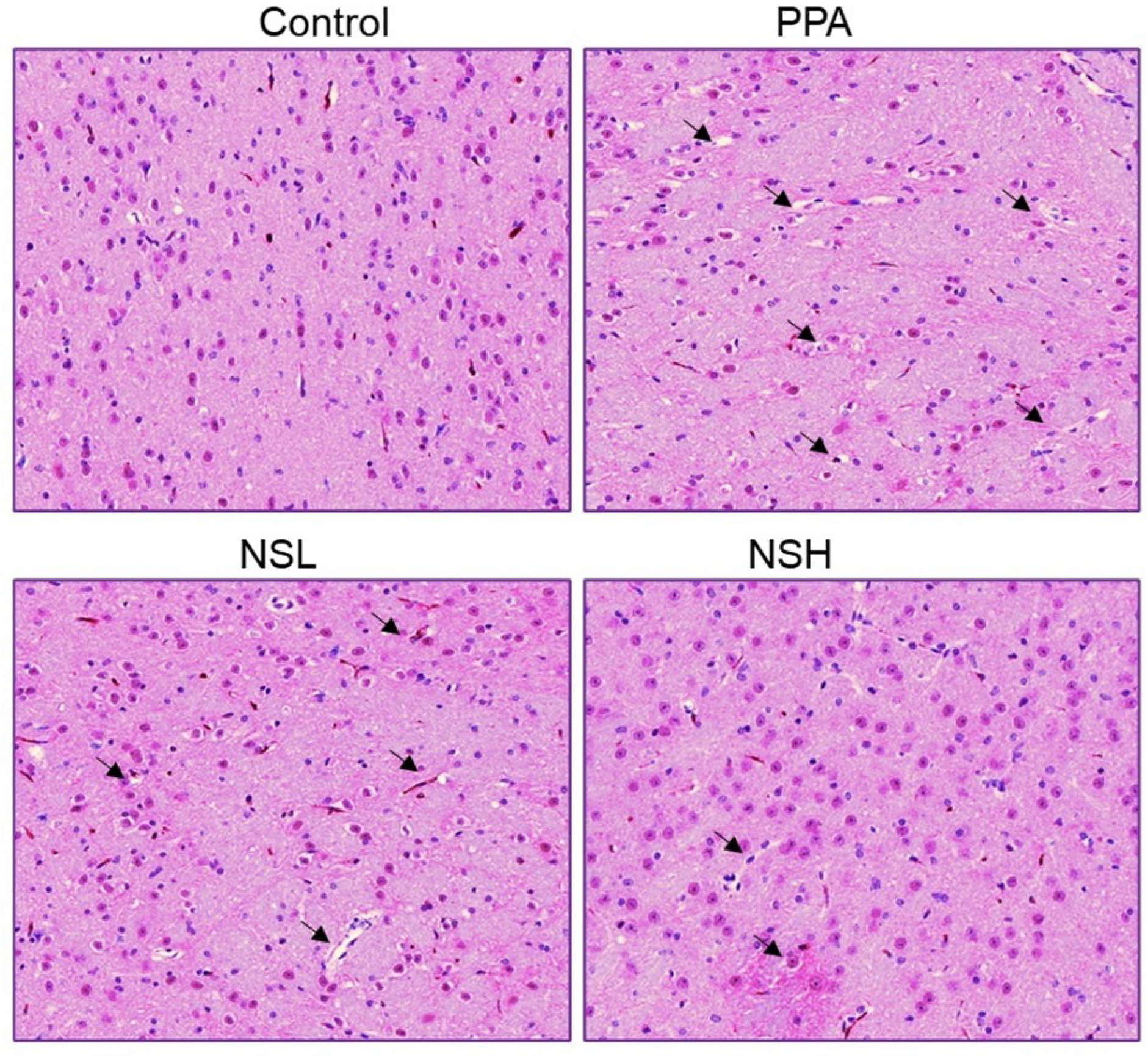
Effect of *Nigella sativa* extract on brain histopathology in autism-spectrum disorder (ASD)-like adult Wistar rats (N=40, n=10/group). H&E-stained brain sections of Control (Group I, NaCl 0.9%); PPA (Group II, Propionic acid); NSL (Group III, *N. sativa* Low-dose); NSH (Group IV, *N. sativa* High-dose). Black arrows depict brain cell damages.

Both *N. sativa*-treated groups (NSL and NSH) showed less neuronal degeneration with 6 ± 2 alterations in the brain tissue of NSL Group, and the NSH Group with 3 ±1 alterations, exhibiting near-normal brain tissue architecture.

## IV. Discussion

Autism Spectrum Disorder (ASD) is a highly complex neurodevelopmental disorder showing a lack of social interaction, communication, and the presence of restricted and repetitive behaviors [29]. Current pharmacological interventions often focus on alleviating symptoms, but they rarely address the underlying neurobiological disruptions or have sustained therapeutic effects [30]. As a result, alternative therapies are increasingly being explored. *N. sativa* (black cumin) has received increased interest in the last few years due to its antioxidant, anti-inflammatory, and neuroprotective properties [11,12]. Morover, one of the active constituents of *N. sativa* Thymoquinone has been shown to exert neuroprotective effects in valproic acid-induced rat model of autism [8].

In this study, we evaluated the effects of *N. sativa* extract in a PPA-induced rat model of ASD. PPA was selected due to its ability to reliably induce core ASD-like features (i.e., social deficits, repetitive behaviors, and anxiety) through mechanisms involving neuroinflammation, oxidative stress, and gut-brain axis disruption [31]. Unlike valproic acid (VPA), which models prenatal risk via epigenetic modulation, PPA mimics postnatal environmental influences and gut microbial metabolites implicated in ASD pathophysiology [32]. Its reproducibility, rapid induction, and clinical relevance make it a rational model for testing therapeutic agents like *N. sativa*. Reciprocal social interaction and stereotypic behavior tests showed statistically different effects between experimental groups, strenghtenening the potential of *N. sativa* as a treatment for ASD. As anticipated, PPA-treated rats exhibited substantial impairments in social behavior and a dramatic enhancement of stereotypic behavior, both of which are hallmark features of ASD. These findings agree with previous reports in which PPA administration leads to ASD-like phenotype including deficits of social interactions and increased repetitive behaviors [33]. Notably, both *N. sativa* treatment groups (NSL and NSH) improved these behaviors significantly. In particular the higher dose (NSH) was quite similar to that of control in terms of social interaction time and frequency. These results are in confirmation with a previous study wherein the phytochemical Thymoquinone attenuates behavioral impairments in a VPA-induced rat model of autism through oxidative stress modulation and inflammation [8]. Further, reports about the neuroprotective properties of the bioactive compounds such as thymoquinone of *N. sativa* have proposed that these compounds may restore normal behavior by influencing neurotransmitter systems and ameliorating neural inflammation [34, 8]. This is in agreement with our present study and strongly suggest an ability of *N. sativa* to affect neural circuits associated with social function and repetitive behaviors.

In a step further, anxiety-like behaviors were more broadly assessed through the Open Field Test (OFT) and Elevated Plus Maze Test (EPMT). These tests are routinely employed to assess exploratory activity and anxiety-related responses in rodent models [35]. Satisfactorily, PPA group of rats displayed significant anxiety, which was evidenced by reduced exploration of the open field and a preference for the enclosed arms of the EPM. These results are consistent with reports from other ASD models in which anxiety is an identified comorbidity [36]. *N. sativa* treatment significantly attenuated these anxiety-like behaviors and the highest dose caused the most significant effects. The NSH group showed enhanced locomotion in both the open field and for the open arms of the EPM, indicating a decrease in anxiety. This aligns with the growing body of evidence supporting the anxiolytic effects of *N. sativa*, which is most likely due to its ability to modulate neurotransmitters involved in mood regulation, such as serotonin and GABA [37].

Interestingly, oxidative stress is a documented pathophysiologic mechanism in ASD and numerous studies have reported increased oxidative stress markers in human patients as well as animal models of ASD [15]. Herein, we report that PPA treatment in rats caused substantial rises in the levels of malondialdehyde (MDA), an index of lipid peroxidation, and in the decreases associated with the activity of glutathione (GSH) and superoxide dismutase (SOD), reflecting an upset of the oxidative-antioxidative system in brain tissues. This observation is also in line with the oxidative stress involvement in neurodevelopmental disorders, such as ASD [38]. Post-treatment with *N. sativa* significantly decreased oxidative stress markers manner the. The reduction in MDA levels, along with the restoration of GSH and SOD activities, suggests that *N. sativa* can effectively counteract oxidative damage in the brain. Thes findings are consistent with a number of reports highlighting the strong *N. sativa*-induced antioxidant activity, most notably of its purported active ingredient, thymoquinone [39, 40, 8], which has been shown to abrogate free radical generation and enhance endogenous antioxidant defence. The normalization of the oxidative stress level in the *N. sativa*-treated groups may account, at least partially, for the behavioral and neurobiological improvements observed in our study.

Importantly, *N. sativa* treatment attenuated the expression of pro-inflammatory cytokines, TNF-α and IL-1β in brain tissues, conversely to what we could have observed in the PPA group of rats. Inflammation in ASD models is often associated or linked to elevated levels of such pro-inflammatory markers, which contribute in the increased neuroinflammation and neurodevelopmental deficits, both characteristics of ASD pathophysiology [41]. Our data agree with the *N. sativa*-mediated anti-inflammatory property shown in other models, mentioning the regulation of pro-inflammatory cytokines and modulation of the immune response [42]. The dramatic drop in the expression of those cytokines in the NSH group, confirms that *N. sativa* influence neuroinflammatory pathways which may lead to additional relief of ASD symptoms (behavioral and physiological).

Moreover, the neurotransmitter dysregulation is an additional hallmark of ASD, specifically in dopamine and serotonin systems [43]. Our findings showed that the brain of PPA group of rats had significantly reduced levels of both dopamine and serotonin, which is consistent with previous studies reporting neurotransmitter imbalances in ASD models [44-46]. Treatment with *N. sativa* resulted in a significant restoration of these neurotransmitters balance which may also contribute to the behavioral improvements observed in our study, as dopamine and serotonin are critical in regulating mood, social behavior, and repetitive movements. The ability of *N. sativa* to normalize these neurotransmitter systems is in line with prior research suggesting that its bioactive compounds, such as thymoquinone, can positively influence neurotransmitter activities in the brain [8, 34, 39, 40].

The decrease in caspase-3 expression in the brain tissue of *N. sativa*-treated groups of rats, compared to the PPA group, further reinforces the neuroprotective action of *N. sativa*, since caspase-3 is a marker of apoptosis [47]. The higher levels of caspase-3 expression in the brain tissue of PPA group of rats are consistent with increased neuronal death observed in multiple ASD models. Treatment of *N. sativa* potently downregulated expression of caspase-3, indicating that the extract possess anti-apoptotic activities. These findings corroborate other research studies that has reported protective effect of *N. sativa* against oxidative stress-mediated neuronal damages, probably by antioxidant and anti-inflammatory activities [48].

Eventually, histopathological analyses of brain tissue showed an extensive neuronal loss and gliosis in the PPA [49]. In contrast, the *N. sativa*-treated groups of rats showed less neuronal degeneration and reduced glial activation, with the highest tested dose (NSH) exhibiting near-normal brain architecture. These results are consistent with previous reports for the neuroprotective effects of *N. sativa* [8, 12, 13], which show the attenuation of oxidative stress and inflammation, ultimately contributing to neuronal survival and healthy brain function.

## Limitations and perspectives of the study

While the current findings demonstrate the neuroprotective potential of *N. sativa* in a PPA-induced model of ASD, several limitations warrant consideration and shall be overcome in a next research project. First, phytochemical profiling of the extract was not performed, and thus, the specific bioactive compounds responsible for the observed effects remain unidentified to date. Second, the study lacks pharmacokinetic data and dose-ranging analyses across broader concentrations, which are essential for optimizing therapeutic efficacy. Third, real-time cerebral imaging was not conducted, which could have provided additional insight into neurofunctional alterations before and after treatment. Furthermore, the molecular signaling pathways underlying the beneficial effects of *N. sativa* were not elucidated; understanding these pathways is crucial for mechanistic validation. The study also did not compare *N. sativa* to other neuroprotective or antioxidant-rich plants, limiting the relative evaluation of its efficacy. Moreover, toxicological assessments, including cytotoxicity, hemocompatibility, and long-term safety at varying doses, were not performed. Lastly, nanoformulation strategies were not explored; such approaches may enhance the bioavailability, brain penetration, and pharmacological stability of *N. sativa* extracts. Future studies incorporating these aspects are programmed in order to establish the therapeutic viability and translational potential of *N. sativa* in ASD and related neurodevelopmental disorders.

## Conclusions

Overall, this study is a step forward the potential of *N. sativa* in ASD management while opening new avenues. This study not only confirms the the capacity of *N. sativa* to act as a natural traditional remedy for neurodisorders, but also report new insights in the reduction of ASD characteristics in an unexplored animal model in which PPA has induced ASD-like symptoms. The findings showed that *N. sativa* extract, exhibits the ability to attenuate behavioural and neurobiological symptoms of ASD observed in a rat model of PPA-induced ASD. The *N. sativa*-mediated enhancements in social behavior, repetitive behaviors, anxiety, and neurotransmitter balance with the decrease of oxidative stress, inflammation, and neuronal loss, indicate the therapeutic prospect of *N*.*sativa* in ASD. Although additional studies are needed to fully describe the pharmaco-toxicological properties of *N. sativa* as well as the underlying molecular mechanisms of this effect in animal models, prior to establish the clinical efficacy of *N. sativa* in humans, our data show that it may provide a potentially effective adjunct therapy in the treatment of core and comorbid symptoms of ASD.

## Acknowledgments

The authors gratefully acknowledge the University of Kashmir, India, as well as King Khalid University, Saudi Arabia, for providing access to core laboratory facilities and technical support essential to the completion of this work. Their infrastructure and collaborative environment significantly contributed to the research outcomes presented in this study.

## Institutional Review Board Statement

Ethical approval for the study was obtained from the Institutional Animal Care and Use Committee (IACUC), and all procedures adhered to national and institutional animal welfare guidelines. In addition, the study was approved by the research ethics committee of University of Kashmir, Hazratbal, Kashmir, India, with approval number UK/2021/AS/10.

## Informed Consent Statement

Not applicable.

## Clinical trial Number

Not applicable.

## Data Availability Statement

The datasets used and/or analyzed during this study are available from the corresponding author(s) upon reasonable request.

## References

1. Hirota T, King BH. Autism spectrum disorder: a review. JAMA. 2023;329(2):157–168.

2. Briot K, Jean F, Jouni A, et al. Social anxiety in children and adolescents with autism spectrum disorders contribute to impairments in social communication and social motivation. Front Psychiatry. 2020;11:710.

3. Lord C, Brugha TS, Charman T, et al. Autism spectrum disorder. Nat Rev Dis Primers. 2020;6(1):1–23.

4. Manoli DS, State MW. Autism spectrum disorder genetics and the search for pathological mechanisms. Am J Psychiatry. 2021;178(1):30–38.

5. Aishworiya R, Valica T, Hagerman R, Restrepo B. An update on psychopharmacological treatment of autism spectrum disorder. Neurotherapeutics. 2023;19(1):248–262.

6. Zhukova MA, Talantseva OI, Logvinenko TI, et al. Complementary and alternative treatments for autism spectrum disorders: a review for parents and clinicians. Clin Psychol Spec Educ. 2020;9(3).

7. Pangrazzi L, Balasco L, Bozzi Y. Natural antioxidants: a novel therapeutic approach to autism spectrum disorders? Antioxidants. 2020;9(12):1186.

8. El-Naggar, M. E., et al. (2021). Thymoquinone ameliorates behavioral impairments in a valproic acid-induced rat model of autism by modulating oxidative stress and inflammation. NeuroToxicology, 85, 1–9.

9. Alharbi SA. Herbal medicine approach and their effectiveness in the management of autism spectrum disorders. Res J Pharm Technol. 2024;17(7):3459–3466.

10. Vandana P, Simkin DR, Hendren RL, Arnold LE. Autism spectrum disorder and complementary-integrative medicine. Child Adolesc Psychiatr Clin N Am. 2023;32(2):469–494.

11. Singh R, Kisku A, Kungumaraj H, et al. Autism spectrum disorders: a recent update on targeting inflammatory pathways with natural anti-inflammatory agents. Biomedicines. 2023;11(1):115.

12. Chaturvedi S, Gupta R, Gupta N, et al. Nigella sativa and its chemical constituents: a promising approach against neurodegenerative disorders. In: Black Seeds (Nigella Sativa). Elsevier; 2022:149–176.

13. Ojueromi OO, Oboh G, Ademosun AO. Black seed (Nigella sativa): a favourable alternative therapy for inflammatory and immune system disorders. Inflammopharmacology. 2022;30(5):1623–1643.

14. Kulsum K, Syahrul S, Hasballah K, Balqis U. Propofol and Nigella sativa L. seeds ethanol extract enhance neuroprotection: a histopathological study in rat models with traumatic brain injury. Indones Biomed J. 2024;16(5):450–458.

15. Bjørklund G, Meguid NA, El-Bana MA, et al. Oxidative stress in autism spectrum disorder. Mol Neurobiol. 2020;57:2314–2332.

16. Willsey HR, Willsey AJ, Wang B, State MW. Genomics, convergent neuroscience and progress in understanding autism spectrum disorder. Nature Reviews Neuroscience. 2022 Jun;23(6):323–41.

17. Gevezova M, Sarafian V, Anderson G, Maes M. Inflammation and mitochondrial dysfunction in autism spectrum disorder. CNS & Neurological Disorders-Drug Targets-CNS & Neurological Disorders). 2020 Jun 1;19(5):320–33.

18. Li Z, Zhu YX, Gu LJ, Cheng Y. Understanding autism spectrum disorders with animal models: applications, insights, and perspectives. Zoological research. 2021 Nov 18;42(6):800.

19. Maryam T, Rana NF, Alshahrani SM, Batool F, Fatima M, Tanweer T, Alrdahe SS, Alanazi YF, Alsharif I, Alaryani FS, Kashif AS, Menaa F. Silymarin Encapsulated Liposomal Formulation: An Effective Treatment Modality against Copper Toxicity Associated Liver Dysfunction and Neurobehavioral Abnormalities in Wistar Rats. Molecules. 2023 Feb 3;28(3):1514.

20. MacFabe DF, Cain DP, Rodriguez-Capote K, Franklin AE, Hoffman JE, Boon F, Taylor AR, Kavaliers M, Ossenkopp KP. Neurobiological effects of intraventricular propionic acid in rats: possible role of short chain fatty acids on the pathogenesis and characteristics of autism spectrum disorders. Behavioural brain research. 2007 Jan 10;176(1):149–69.

21. Hobbenaghi R, Javanbakht J, Sadeghzadeh SH, et al. Neuroprotective effects of Nigella sativa extract on cell death in hippocampal neurons following experimental global cerebral ischemia-reperfusion injury in rats. J Neurol Sci. 2014;337(1–2):74–79.

22. Moy SS, Nadler JJ, Young NB, Nonneman RJ, Segall SK, Andrade GM, Crawley JN, Magnuson TR. Social approach and repetitive behavior in eleven inbred mouse strains. Behavioural brain research. 2008 Aug 5;191(1):118–29.

23. Pellow S, Chopin P, File SE, Briley M. Validation of open: closed arm entries in an elevated plus-maze as a measure of anxiety in the rat. Journal of neuroscience methods. 1985 Aug 1;14(3):149–67.

24. Smolinsky AN, Bergner CL, LaPorte JL, Kalueff AV. Analysis of grooming behavior and its utility in studying animal stress, anxiety, and depression. Mood and anxiety related phenotypes in mice: Characterization using behavioral tests. 2009:21–36.

25. ZA P. Estimation of product of lipid peroxidation (malonyldialdehyde) in biochemical systems. Anal Biochem. 1966;16:359–364.

26. Ellman GL. Tissue sulfhydryl groups. Arch Biochem Biophys. 1959;82(1):70–77.

27. Marklund SL, Adolfsson R, Gottfries CG, Winblad B. Superoxide dismutase isoenzymes in normal brains and in brains from patients with dementia of Alzheimer type. J Neurol Sci. 1985;67(3):319–325.

28. Ullah A, Razzaq A, Alfaifi MY, Elbehairi SEI, Menaa F, Ullah N, Shehzadi S, Nawaz T, Iqbal H. Sanguinarine Attenuates Lung Cancer Progression via Oxidative Stress-induced Cell Apoptosis. Curr Mol Pharmacol. 2024;17:e18761429269383.

29. Genovese A, Butler MG. Clinical assessment, genetics, and treatment approaches in autism spectrum disorder (ASD). Int J Mol Sci. 2020;21(13):4726.

30. Joon P, Kumar A, Parle M. The resemblance between the propionic acid rodent model of autism and autism spectrum disorder. J Psych Neurochem Res. 2023;1(3):1–8.

31. Sharma AR, Batra G, Saini L, Sharma S, Mishra A, Singla R, Singh A, Singh RS, Jain A, Bansal S, Modi M. Valproic acid and propionic acid modulated mechanical pathways associated with autism spectrum disorder at prenatal and neonatal exposure. CNS & Neurological Disorders-Drug Targets-CNS & Neurological Disorders). 2022 Jun 1;21(5):399–408.

32. Sahin K, Orhan C, Karatoprak S, Tuzcu M, Deeh PB, Ozercan IH, Sahin N, Bozoglan MY, Sylla S, Ojalvo SP, Komorowski JR. Therapeutic effects of a novel form of biotin on propionic acid-induced autistic features in rats. Nutrients. 2022 Mar 17;14(6):1280.

33. Bhandari R, Sodhi RK, Kuhad A. Neuropsychopathology And Neurobehavioral Characteristics of PPA-Induced Autism Like Rat Model and Its Correlation with Gut-Brain Dysbiosis Occurring in Autism Spectrum Disorder. InAnimal Models for Neurological Disorders 2021 Dec 23 (pp. 153–181). Bentham Science Publishers.

34. Sedaghat R, Roghani M, Khalili M. Neuroprotective effect of thymoquinone, the Nigella sativa bioactive compound, in 6-hydroxydopamine-induced hemi-parkinsonian rat model. Iran J Pharm Res. 2014;13(1):227.

35. de Figueiredo Cerqueira MM, Castro MM, Vieira AA, Kurosawa JA, do Amaral Junior FL, de Siqueira FD, Sosthenes MC. Comparative analysis between Open Field and Elevated Plus Maze tests as a method for evaluating anxiety-like behavior in mice. Heliyon. 2023 Apr 1;9(4).

36. Benitah KC, Kavaliers M, Ossenkopp KP. The enteric metabolite, propionic acid, impairs social behavior and increases anxiety in a rodent ASD model: examining sex differences and the influence of the estrous cycle. Pharmacology Biochemistry and Behavior. 2023 Oct 1;231:173630.

37. Bano F, Ahmed A, Parveen T, Haider S. Anxiolytic and hyperlocomotive effects of aqueous extract of Nigella sativa L.seeds in rats. Pak J Pharm Sci. 2014;27.

38. González-Fraguela ME, Hung ML, Vera H, et al. Oxidative stress markers in children with autism spectrum disorders. Br J Med Med Res. 2013;3(2):307–317.

39. El-Naggar T, Gómez-Serranillos MP, Palomino OM, et al. Nigella sativa L. seed extract modulates the neurotransmitter amino acids release in cultured neurons in vitro. Biomed Res Int. 2010;2010(1):398312.

40. Ismail M, Al-Naqeep G, Chan KW. Nigella sativa thymoquinone-rich fraction greatly improves plasma antioxidant capacity and expression of antioxidant genes in hypercholesterolemic rats. Free Radic Biol Med. 2010;48(5):664–672.

41. Gholamnezhad Z, Keyhanmanesh R, Boskabady MH. Anti-inflammatory, antioxidant, and immunomodulatory aspects of Nigella sativa for its preventive and bronchodilatory effects on obstructive respiratory diseases: a review of basic and clinical evidence. J Funct Foods. 2015;17:910–927.

42. Young AM, Chakrabarti B, Roberts D, et al. From molecules to neural morphology: understanding neuroinflammation in autism spectrum condition. Mol Autism. 2016;7:1–8.

43. Blum K, Bowirrat A, Sunder K, et al. Dopamine dysregulation in reward and autism spectrum disorder. Brain Sci. 2024;14(7):733.

44. Cetin FH, Tunca H, Guney E, Iseri E. Neurotransmitter systems in autism spectrum disorder. Autism Spectrum Disorder—Recent Advances. 2015 Apr 2:15–30.

45. Al-Radadi NS, Al-Bishri WM, Salem NA, ElShebiney SA. Plant-mediated green synthesis of gold nanoparticles using an aqueous extract of Passiflora ligularis, optimization, characterizations, and their neuroprotective effect on propionic acid-induced autism in Wistar rats. Saudi Pharm J. 2024;32(2):101921.

46. Fouad IA, Sharaf NM, Abdelghany RM, El Sayed NS. Neuromodulatory effect of thymoquinone in attenuating glutamate-mediated neurotoxicity targeting the amyloidogenic and apoptotic pathways. Front Neurol. 2018;9:236.

47. Chen DL, Engle JT, Griffin EA, et al. Imaging caspase-3 activation as a marker of apoptosis-targeted treatment response in cancer. Mol Imaging Biol. 2015;17:384–393.

48. Madkour DA, Ahmed MM, Orabi SH, et al. Nigella sativa oil protects against emamectin benzoate-induced neurotoxicity in rats. Environ Toxicol. 2021;36(8):1521–1535.

49. Livingston LA, Happé F. Conceptualising compensation in neurodevelopmental disorders: reflections from autism spectrum disorder. Neurosci Biobehav Rev. 2017;80:729–742.

